# Invasion of malaria mosquitoes natural population by parasitic nematodes Dirofilaria along Ob River basin in Western Siberia

**DOI:** 10.1101/2022.04.04.487071

**Authors:** Vladimir A. Burlak, Valentina S. Fedorova, Gleb N. Artemov

## Abstract

Dirofilariasis – is a dangerous disease that affects carnivores, felines, and humans. It is caused by parasitic nematodes *Dirofilaria repens* and *D. immitis*. These parasites are carried by blood–sucking mosquitoes. In recent decades the habitat of Dirofilaria in Europe has been expanding dramatically. In the present study, we investigated how malaria mosquitoes had been infected by Dirofilaria in Western Siberia (Russia) in the range between 50° 48’ N (Labytnangi) and 66° 39’ N (Kurota) along the Ob River. The extensiveness of parasite infection varied between 0.4 % and 5.7 %, in three species of malaria mosquitoes: *Anopheles beklemishevi, An. daciae, An. messeae*, which all are showed effective vectors of *D. repens*. The results demonstrate the significant role of malaria mosquitoes for dirofilariasis transmission in severe climate conditions of Western Siberia.

---

Dirofilariasis is a dangerous disease of the canine and feline animals as well as humans, which is caused by heterogenous helminths – Dirofilaria. There are two species of Dirofilaria: *Dirofilaria repens* causes subcutaneous form of the disease and *D. immitis* («heartworm») – more severe pulmonary form. Human dirofilariasis is associated with both forms and recently has been registered in many countries, although the human is an aberrant host for Dirofilaria (Capelli et al. 2018).

Last few decades, the rate of dirofilariasis case detection was high in the northern border, which can be explained by dog migrations as well as global warming. (Genchi et al. 2011; Okulewicz 2017). In Europe, the cases of dirofilariasis in dogs and humans had been detected not only in the Mediterranean and Black Sea region but also in Baltic and Nordic countries (Tiškina and Jokelainen 2017; Sabūnas et al. 2019).

The geographic distribution of autochthonous dirofilariasis is associated with the adaptation of nematodes to new climatic conditions and new intermediate hosts – bloodsucking mosquitoes, that are the vectors of the parasites. At least in southern Russia, where it has a high level of population of non–malaria mosquitoes, these genera seems to be the major vector of dirofilariasis. In the northern region, malaria mosquitoes are the major vectors (Rakova et al., 2012, Kronefeld et al, 2014). Russia is the country with the highest rate of human dirofilariasis infection counting for one–third of the total cases in Europe. (González–Miguel et al. 2020). Due to its vast territory, some regions have not been or had only been partially studied for the invasiveness of the disease and the vector species.

The Western–Siberian plain is bordered by the Ural Mountains on the west, Central Siberian Plateau and Yenisei River on the east, Kazakh Uplands on the south, and the Kara Sea on the north. The biggest Ob River flows through the plain from the Altay Mountain on the southern–east to the Gulf of Ob on the northern–west. The basin of the Ob, including the biggest swamp, created the condition for the habitation of various blood–sucking mosquito species, potential vectors of Dirofilariasis. The first case of indigenous transmission of dirofilariasis was recorded in Western Siberia two decades ago – in Novosibirsk and Omsk cities in 1998–1999 yy (Tolkunova et al., 2003; Kronefeld, 2014) and Tomsk and Kolpashevo cities in 2016 (Poltoratskaya et al., 2018). But up till today, the vector–borne parasite invasion in Western Siberia has not been investigated resulting in restrictions on the investigation of the capacity of both dirofilariasis and its host spread in severe conditions such as the northern regions. Non– malaria potential vectors were investigated recently in the border of the Tomsk region (Poltoratskaya et al., 2021), but the role of malaria mosquito fauna plays in dirofilariasis transmission is still underestimated.

There are four species of malaria mosquitoes inhabiting in Western Siberia – *An. messeae, An. daciae, An. beklemishevi* (Artemov et al., 2018; 2021) and *An. claviger* (Poltoratskaya et al., 2021). In this study, we investigated malaria mosquitoes from 20 natural population along Ob River 1) to estimate the rate of mosquito invasion in Western Siberia; 2) to define the northern and the southern borders of dirofilariasis 3) to discover potential and effective vectors within species of malaria mosquitoes and the species parasitic nematodes on the Western Siberia plane.

Mosquitoes were collected in stables, using an aspirator, from May to August, in 2020 and 2021 yy. Each mosquito was dissected on the slide with a drop of PBS solution. Malpighian tubules and proboscis were left on the slide, washed with a new portion of PBS, and covered by coverslip 18×18 mm, where the body was fixed separately in 96% ethanol. Organs were observed microscopically using 5×, 10× and 40× objectives for parasitic helminthes and for measuring the developmental stages (Montarsi et al., 2015). In the case of invasion, the coverslip was removed and the organ remains washed in the 0,5 ml tube with 96% ethanol and stored in –20 °C conditions.

DNA extraction process from Malpighian tubules and proboscis of mosquitoes with nematodes inside was performed according to the protocol (Debrunner–Vossbrinck B.A. et al, 2006) with several modifications. Before the extraction procedure, the material was removed from ethanol, dried at room temperature, and homogenized in 50 μl STE buffer (100 mM NaCl; 10 mM Tris–HCl, pH 8.0; 1 mM EDTA). The homogenate was incubated at 95 °C for 5 min, immediately frozen at –20 °C, and thawed at room temperature to release the DNA from the tissues after 5 min centrifugation at 10000 g. The supernatant with DNA was transferred into the clean 1.5 ml tube and stored at –20° C to be used for PCR reaction.

Parasites species identification was performed by PCR analysis with primers designed for molecular diagnostics of filariasis in dogs, based on the difference in the sequences of COI gene of the mitochondrial DNA (Rishniw M. et al, 2006). Malaria mosquito species identification was performed using PCR and PCR–RFLP analysis of internal transcribed spacer 2 of ribosomal genes cluster (ITS2, Vaulin, et al., 2018, Artemov et al., 2021).

Twenty geographical locations from 50 to 66 northern latitudes have been observed using straightforward analysis of 4077 mosquitoes and 53 from which were infected by Dirofilaria (Table 1). Malaria mosquitoes from the Maculipennis subgroup were found in all observed geographical points with different numbers, depending on the season, time, population phase, and latitude. *An. claviger* has never been detected in stables. The measured area of Dirofilaria infection in malaria mosquitoes was from 62° 32’ N – 62° 57’ N between Priob’e and Peregryobnoye in the north and 52° N between Katunskiy and Sovetskoye in the south (Figure 1). The rate of infection of the parasite from the north to south as a tendency varied from 0.4 % in Shapsha to 5.7 % in Teguldet.

**Table 1.** Extensiveness of invasion and the species of malaria mosquitoes invaded with *D. immitis* and *D. repens* in the Ob River settlements

| № | Settlement | Coordinates | Date | Extensiveness of invasion, % (N) | N<sup>+</sup> | Parasite | DS | Malaria mosquito host |
| --- | --- | --- | --- | --- | --- | --- | --- | --- |
| 1 | Labytnangi, YNAO | 66°39'18"N 66°25'06"E | 09 Aug 2021 | 0 (9) | 0 | – | – | – |
| 2 | Shuryshkary, YNAO | 65°54'05"N 65°20'52"E | 08 Aug 2021 | 0 (126) | 0 | – | – | – |
| 3 | Berezovo, KhMAO | 63°56'24"N 65°01'35"E | 05 Aug 2020<br>30 June 2021 | 0 (351) | 0 | – | – | – |
| 4 | Peregryobnoye, KhMAO | 62°57'48"N 65°05'55"E | 26 June 2021 | 0 (420) | 0 | – | – | – |
| 5 | Priob'e, KhMAO | 62°32'38"N 65°38'18"E | 27 June 2021 | 1.1 ± 0.5 (370) | 4 | *D. repens* | L1 | *An. messeae* |
| 6 | Yaguryakh, KhMAO | 61°15'17"N 67°40'01"E | 27 June 2021 | 0 (313) | 0 | – | – | – |
| 7 | Shapsha, KhMAO | 61°05'24"N 69°27'49"E | 17–28 July 2021 | 0.4 ± 0.3 (510) | 2 | *D. repens* | L1 | *An. messeae* |
| 8 | Strezhevoy, TO | 60°42'59"N 77°33'33"E | 23 July 2020 | 1.0 ± 0.7 (206) | 1 | *D. repens* | L2 | *An. messeae* |
|  |  |  |  |  | 1 | *D. repens* | – | *An. beklemishevi* |
| 9 | Bolshaya Griva, TO | 58°55'33"N 80°14'32"E | 16 July 2020 | 1.1 ± 1.1 (95) | 1 | *D. repens* | L3 | *An. beklemishevi* |
| 10 | Klukvinka, TO | 58°32'26"N 85°51'22"E | 28 May 2020 | 2.6 ± 1.3 (154) | 3 | *D. repens* | L1 | *An. beklemishevi* |
| 11 | Kolpashevo agglomeration<br>(villages: Novoselovo,<br>Bolshaya Sarovka, Togur),<br>TO | 58°19'27"N 82°55'10"E | 02 July 2020 | 3.9 ± 0.8 (540) | 11 | *D. repens* | L1–2 | *An. messeae* |
|  |  |  |  |  | 3 | *D. repens* | L1–2 | *An. daciae* |
|  |  |  |  |  | 3 | *D. repens* | L2 | *An. beklemishevi* |
|  |  |  |  |  | 1 | *D. immitis* | L2 | *An. messeae* |
|  |  |  |  |  | 1 | *D. repens, D. immitis\** | L1, L2 | *An. daciae* |
|  |  |  |  |  | 1 | *D. repens, D. immitis\** | L2 | *An. beklemishevi* |
| 12 | Teguldet, TO | 57°18'52"N 88°09'22"E | 21 May 2020 | 5.7 ± 2.2 (106) | 5 | *D. repens* | L1 | *An. messeae* |
|  |  |  |  |  | 1 | *D. repens* | L1 | *An. daciae* |
| 13 | Kolarovo, TO | 56°19'56"N 84°57'48"E | 29 Aug 2020 | 3.4 ± 1.9 (89) | 4 | *D. repens* | L1–3 | *An. daciae* |
| 14 | Roshinskiy, NO | 54°37'59"N 83°13'02"E | 25 July 2021 | 1.4 ± 0.7 (287) | 3 | *D. repens* | L1 | *An. daciae* |
|  |  |  |  |  | 1 | *D. repens* | L1 | *An. beklemishevi* |
| 15 | Novoaltaysk, AK | 53°27'19"N 83°54'41"E | 17 Aug 2020 | 4.7 ± 2.6 (64) | 2 | *D. immitis* | L3 | *An. daciae* |
| 16 | Katunskiy, AK | 52°24'35"N 85°07'18"E | 04 Aug 2021 | 3.5 ± 1.9 (85) | 1 | *D. repens, D. immitis\** | L3 | *An. messeae–daciae* hybrid |
|  |  |  |  |  | 2 | *D. repens* | L1,2 | *An. messeae–daciae* hybrid |
| 17 | Sovetskoye, AK | 52°17'12"N 85°25'01"E | 04 Aug 2021 | 0 (45) | 0 | – | – | – |
| 18 | Choya, RA | 52°01'47"N 86°32'22"E | 05 Aug 2021 | 0 (84) | 0 | – | – | – |
| 19 | Ozernoje, RA | 51°49'02"N 85°48'04"E | 19 Aug 2020<br>06 Aug 2021 | 0 (181) | 0 | – | – | – |
| 20 | Kurota, RA | 50°48'39"N 85°58'15"E | 07 Aug 2021 | 0 (42) | 0 | – | – | – |
| Total |  |  |  | 53 from 4077 | 51 | – |  |  |

YNAO – Yamal–Nenets Autonomous Okrug, KhMAO – Khanty–Mansi Autonomous Okrug, TO – Tomsk Oblast, NO – Novosibirsk Oblast, AK – Altay Kray, RA – Republic of Altay. N – Number of individual mosquitoes studied; N<sup>+</sup> – number of individuals, invaded by dirofilariae and for which host–parasite pairs were defined, \* – mix–invasion; DS – microfilariae development stag

**Figure 1.**
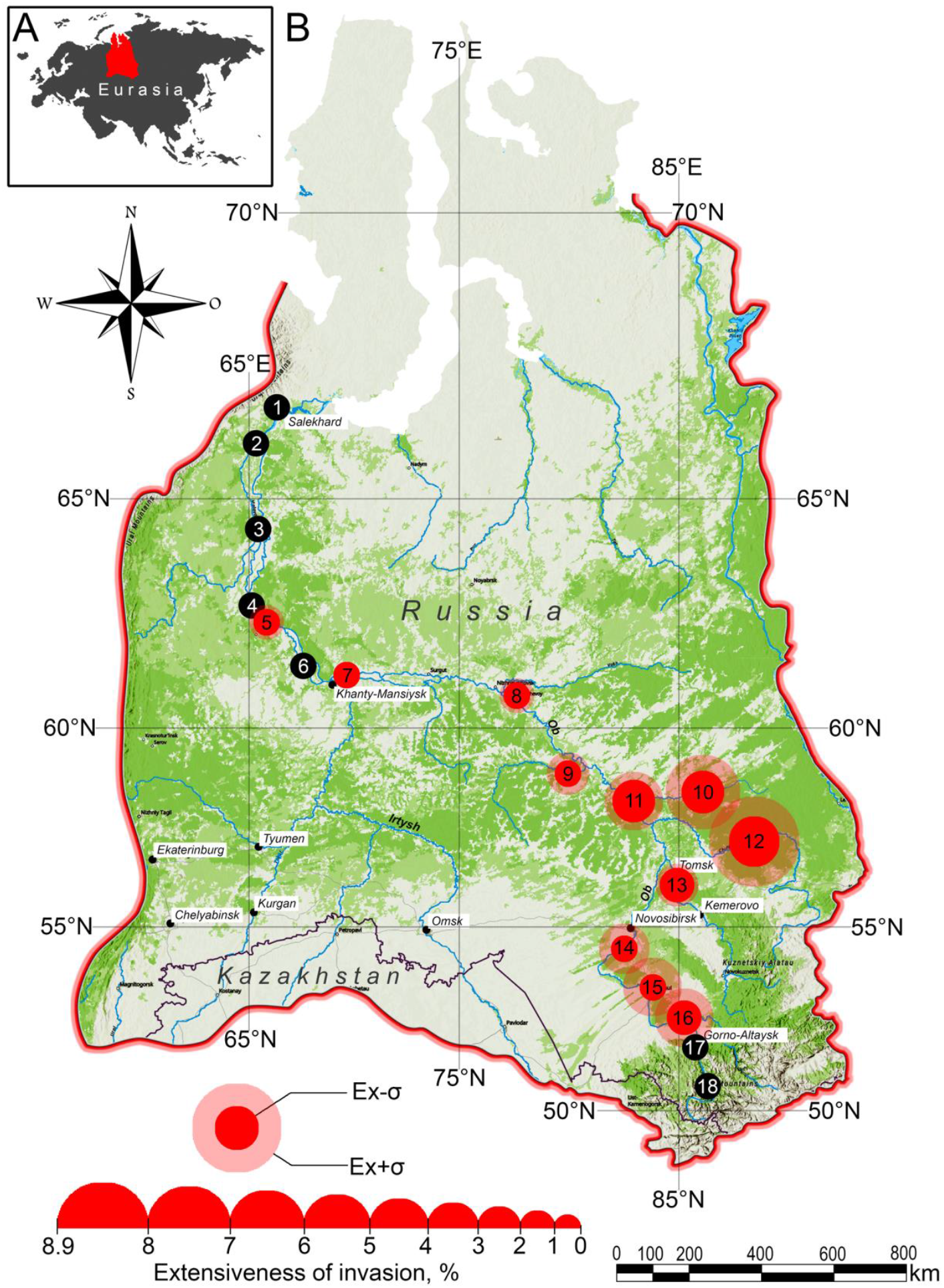
Location of Western–Siberia Plane in Eurasia (A) and the distribution of *D. repens* and *D. immitis* in malaria mosquitoes along Ob River basin (B). Black circles indicate observed settlements with no invaded malaria mosquitoes. The size of red (or pink) circles reflects the extensiveness of invasion accordingly the scale bar on the bottom. Ex – extensiveness of invasion.

In Labytnangi we have identified all three species – *An. beklemishevi, An. messeae*, and *An. daciae*. Thus, we have changed the northernmost range of the area of the malaria mosquitoes *An. beklemishevi, An. messeae* and *An. daciae* distribution in West Siberia from 63° 56’ N in 1974 (Berezovo, Stegniy et al., 1976) to 66° 39’ N (Labytnangi). The detection of *An. daciae* in Western Siberia for the first time on the northernmost border close to the Arctic Circle is evidence of its resistance to the severe conditions against our previous assumption of it being a thermophilic species.

The area of the spread of Dirofilaria corresponds to temperature +14 °С in July, in isotherm zone observed from 2009 to 2019 (Kondrashin et al, 2020), and was confined within the region of 63° 56’ N (Berezovo) and at 51° 24’ N (Chemal, in Altay mountains). We have not detected mosquitoes in Berezovo in 2020, 2021, and Peregrebnoye in 2021, probably due to transport restrictions, making the area only reachable by air or water. Priob’e is the nearest point to Berezovo in the north, which is situated on the highway.

The invasiveness is lower than the epidemically significant level (Sergiev et al., 2014), despite local cases of dirofilariasis detecting in Kolpashevo (Poltoratskaya et al., 2018) and Priob’e (social media information).

*An. messeae, An. daciae* and *An. beklemishevi* were identified as the vectors of Dirofilaria (Table 1). In the area between 58° 19’ N (Kolpashevo ag.) and 57° 18’ N (Teguldet) where are all three species of malaria mosquitoes were found. But in northerner Kolpashevo ag. – only two species were found, *An. messeae* and *An. beklemishevi*. In the southerner Teguldet – both *An. beklemishevi* and *An. daciae* appeared as potential vectors. There were three specimens from Katunskiy village that belonged to the hybrid form of *An. messeae* and *An. daciae*.

*An. messeae* s. l. and *An. daciae* were reported as the vectors for dirofilariasis in Europe earlier (Shaikevich et al., 2019, Ionica et al., 2017), what’s more *An. beklemishevi* was detected for being the vector of *D. repens* for the first time. Three hybrid specimens detected in Novoaltaysk are not unique cases, as one mosquito was found recently in Egoryevsk (Naumenko et al., 2020), but we have found *messeae–daciae* hybrid for the first time in the present study.

Both parasite species, *D. repens* and *D. immitis* were found in malaria mosquitoes. *D. repens* was the predominant species, whereas *D. immitis* was identified only in six out of 51 samples. Notably, half of these samples were detected with both parasites. It was represented by the set of Dirofilaria on different or same developmental stages in Malpighian tubules within the same species of mosquitoes.

The spread of *D. immitis* is confined within 58° 19’ N (Kolpashevo ag.), where two cases of this parasite infection in mosquitos were found. Both cases were detected in the south, Altay valley (Novoaltaysk and Katunskiy), one along, the other with *D. repens* (mix–invasion).

*D. repens* was the dominant parasite in the observed mosquito population. This can be explained by the low level of appearance of *D. immitis* in its definitive host in the Western Siberia region. The prevalence of *D. immitis* in infections containing both parasites has no simple explanation.

We have found Dirofilaria in every developmental stage of mosquitoes. L1 was predominant and observed in ∼60 % of mosquitoes, whereas L2 appeared in ∼30 % and L3 in ∼10 % of the cases (Table 1). Infection of Dirofilaria was detected in Bolshaya Griva and this settlement is the northernmost point of the spread of the disease. The infective stages of Dirofilaria were observed in malaria mosquitoes, in Western Siberia, corresponding to 58° 55’ northern latitude. Two samples of *D. repens* were found in the proboscis of the female *An. beklemishevi*. The infective stage of Dirofilaria was observed also in *An. daciae* from Kolarovo village (*D. repens*) and Novoaltaysk town (*D. immitis*). One remarkable case of double parasite infection in the hybrid of *An. messeae* and *An. daciae* was studied in Katunskoye. The Dirofilaria developed up to the L3 stage and were observed in Malpighian tubules, with no other parasite in proboscis and thorax. Nevertheless, PCR was explained by double infection as a result of the spread of both D. repens and D. immitis.

Out of 50 investigated mosquitoes affected by *D. repens*, 45.09 ± 6.97 % belonged to *An. messeae*, whereas *An. daciae* and *An. beklemishevi* only accounted for 27.45± 6.24 % and 19.61± 5.97 % respectively. All three species and even a hybrid between *An. messeae* and *An. daciae* were the hosts for *D. immitis*, but half of them carried both parasites. *D. immitis* alone was found in *An. messeae* from Kolpashevo ag. population and *An. daciae* in Novoaltaysk.

## Conclusion

The present study has demonstrated that dirofilariasis is a widespread disease in Eurasia and even spread to severe climatic zones in Siberia. The possibility of Dirofilaria spread is very high, as they have occupied new hosts, at least, *An. beklemishevi* was shown as a new vector for *D. repens*. Thus, tree Siberian malaria mosquitoes species, belonging to the Maculipennis subgroup, played an important role as vectors of dirofilariasis in Western Siberia.

## Acknowledgements

The study was supported by a grant from the Russian Science Foundation (Project No. 20–74–10040)

## Notes

### Competing Interest Statement

The authors have declared no competing interest.

